# Condensin chromatin association is regulated by SMC head and hinge engagement and phosphorylation

**DOI:** 10.1101/2025.11.17.688780

**Authors:** Catrina A. Miles, Miao Yu, Nicola E. Minchell, Stephanie A. Schalbetter, Antony W. Oliver, Steve M. Sweet, Jonathan Baxter

## Abstract

The SMC condensin complex is essential for normal mitotic chromosome structure in eukaryotes. Here, we analyze how ATP binding and hydrolysis, SMC hinge stability, and condensin subunit phosphorylation influence condensin chromatin association in budding yeast. We show that mutations predicted to destabilize ATP binding and head-to-head engagement impair condensin chromatin association as assayed by ChIP. In contrast, mutations that inactivate the complex by destabilizing the hinge domain interaction, enhance chromatin association at known condensin loading sites. We find that condensin phosphorylation is enriched in enzymatic states associated with elevated chromatin binding. Moreover, phosphorylation and Aurora/Ipl1 kinase activity—but not Cdc5—are required to maintain condensin association with mitotic chromatin in early metaphase.

## Introduction

Structural maintenance of chromosome (SMC) complexes are universally required to organize replicated chromosomes and enable faithful segregation (Hirano, 2012). Eukaryotes express several distinct SMC complexes, each playing a crucial role in organizing chromosomes to support accurate gene expression and inheritance. The condensin SMC complex is closely associated with mitotic chromosome compaction, where it is required to promote chromosome loop arrays in mitosis (Gibcus *et al*, 2018) The SMC cohesin complex promotes sister chromatid cohesion between replicated chromosomes (Uhlmann, 2016) whilst also organizing chromosome into distinct functional domains by chromosome looping (Rao *et al*, 2017; Schalbetter *et al*, 2017; Schwarzer *et al*, 2017). SMC5/6 is associated with genome stability and DNA repair (Aragon, 2018). *In vitro* cohesin, condensin, SMC5/6 and bacterial SMC complexes directly generate *cis-*DNA loops by loop extrusion (Davidson *et al*, 2019; Ganji *et al*, 2018; Pradhan *et al*, 2025; Pradhan *et al*, 2023; Roisne-Hamelin *et al*, 2025). In this process DNA bound SMC complexes use ATP hydrolysis to bind and translocate DNA (Burmann & Lowe, 2023).

SMC complex activities are generated by a conserved architecture formed by a dimer of SMC proteins in complex with multiple non-SMC subunits (Burmann & Lowe, 2023). Each SMC protein contains an ABC-type ATPase at its ‘head’ domain, linked via a 50nm intramolecular coiled-coil to the major dimerization interface found at its ‘hinge’. When dimerized the SMC proteins are also linked by a non-SMC kleisin factor which binds through its N terminus to the coiled coil region proximal to the head domain of the **γ** SMC and the head domain of the **κ** SMC. The kleisins of cohesin and condensin also interact with a specific subset of HEAT domain proteins collectively referred to as HAWK proteins (Wells *et al*, 2017). Condensin is formed through hetero-dimerization of the SMC proteins Smc2 and Smc4, which in budding yeast are bridged by the kleisin protein Brn1. In addition, Brn1 binds two HAWK proteins Ycg1 and Ycs4 to generate a pentameric complex (Lee *et al*, 2020). All five budding yeast condensin factors are essential for condensin function (Bhalla et al., 2002; Freeman et al., 2000; Lavoie et al., 2002; Ouspenski et al., 2000).

Understanding which enzymatic steps regulate the engagement of SMC complexes with chromatin and which are required for subsequent chromosome organizing action such as the generation of *cis-*loops is a key aspect of understanding how SMCs reconfigure chromosomes (Burmann & Lowe, 2023).

In budding yeast, mutation of any condensin subunit inhibits chromatin association (Lavoie *et al*, 2002), suggesting that the interaction of all subunits is required for stable chromatin binding. SMC complexes have several direct DNA binding surfaces. For condensin, the non-SMC subunits Ycg1 and Brn1 of the budding yeast complex mediate direct DNA binding activity (Kschonsak *et al*, 2017; Piazza *et al*, 2014). Evidence for the head domains directly interacting with DNA has been reported for the *E. coli* SMC MukB and RAD50 (Rojowska *et al*, 2014; Woo *et al*, 2009). DNA binding is also observed for the hinges of *Bs*SMC, cohesin, fission yeast and mouse condensin and Smc5/6 (Akai *et al*, 2011; Alt *et al*, 2017; Griese *et al*, 2010; Hirano & Hirano, 2002, 2006). Both *in vivo* and *in vitro,* ATP hydrolysis by SMC complexes is required for “topological entrapment” of DNA – a stable state in which the complex entraps DNA duplex without requiring direct electrostatic interactions with DNA (Murayama & Uhlmann, 2015; Wilhelm *et al*, 2015). Studies on the *Bs*SMC complex have indicated that stable DNA binding occurs through the combined action of the hinge and head domains utilizing both ATP hydrolysis-dependent and -independent pathways (Hirano & Hirano, 2006; Minnen *et al*, 2016). Current *in vitro* and *in vivo* analyses indicate that the non-SMC subunits make initial contact with DNA, followed by a clamping step triggered by ATP binding and the engagement the SMC heads. ATP hydrolysis is then required to generate the topologically entrapped state (Burmann *et al*, 2025; Burmann & Lowe, 2023).

Mitotic condensin phosphorylation has also been reported to regulate condensin’s chromatin association, via CDK, Aurora and Polo type kinase activity (Giet & Glover, 2001; Kimura *et al*, 1998; Lavoie *et al*, 2004; Lipp *et al*, 2007; Nakazawa *et al*, 2011; Robellet *et al*, 2015; St-Pierre *et al*, 2009; Sutani *et al*, 1999; Tada *et al*, 2011; Thadani *et al*, 2018). Phosphorylation of yeast condensin subunits is focused on the flexible regions of non-SMC subunits (by Ipl1/Cdc5 kinases) and the yeast specific SMC4 N terminal region (CDK) (Robellet *et al*., 2015; St-Pierre *et al*., 2009; Thadani *et al*., 2018). In extracts and *in vitro* the non-SMC flexible regions can inhibit condensin complex function, likely by inhibiting the enzymatic cycle of the complex (Cutts *et al*, 2025; Tane *et al*, 2022). Phosphorylation of these peptide regions has been proposed to alleviate enzymatic inhibition (Cutts *et al*., 2025; Tane *et al*., 2022). These findings predict *in vivo* connections between the enzymatic cycle of condensin, condensin subunit phosphorylation and condensin complex chromatin association which we test here in the budding yeast *Saccharomyces cerevisiae*.

## Results

To examine how condensin’s chromatin association relates to its ATP-binding and hydrolysis cycle, we generated a series of SMC2 mutants carrying single amino acid substitutions (Figure 1A). Several of these mutants were based on well characterized substitutions in bacterial and eukaryotic SMCs that either block ATP binding (the Walker A mutant K38I or Walker B mutant D1112A) (Arumugam *et al*, 2003; Hirano *et al*, 2001; Hirano & Hirano, 1998), reduce ATP hydrolysis (E1113Q) (Arumugam *et al*., 2003; Hirano *et al*., 2001) or alleviate DNA stimulated ATP hydrolysis (R58A) (Lammens *et al*, 2004). We also designed mutations that specifically disrupt one or other of the two hinge hetero-dimerization interfaces of Smc2 (L567K and L665R, Figure 1B), similar to those previously reported for cohesin (Mishra *et al*, 2010). Since all mutations were predicted to lead to loss of condensin’s essential function, we assessed the phenotypes of these mutants using a conditional protein exchange mechanism (Figure 1C). In this system, each of the *SMC2* mutant genes was placed under the control of a galactose inducible promoter (*GAL1p*) and integrated into a strain where a degron cassette has been added to endogenous *SMC2* and transcription controlled through tetR/tetO (Tanaka & Diffley, 2002). Under permissive conditions (glucose or raffinose at 25°C), the degron fused wild-type gene is expressed. After shift to restrictive conditions (+DOX, galactose at 37°C), the degron fused Smc2 is degraded and the *GAL1p* regulated *SMC2* expressed (Figure 1C).

**Figure 1.**
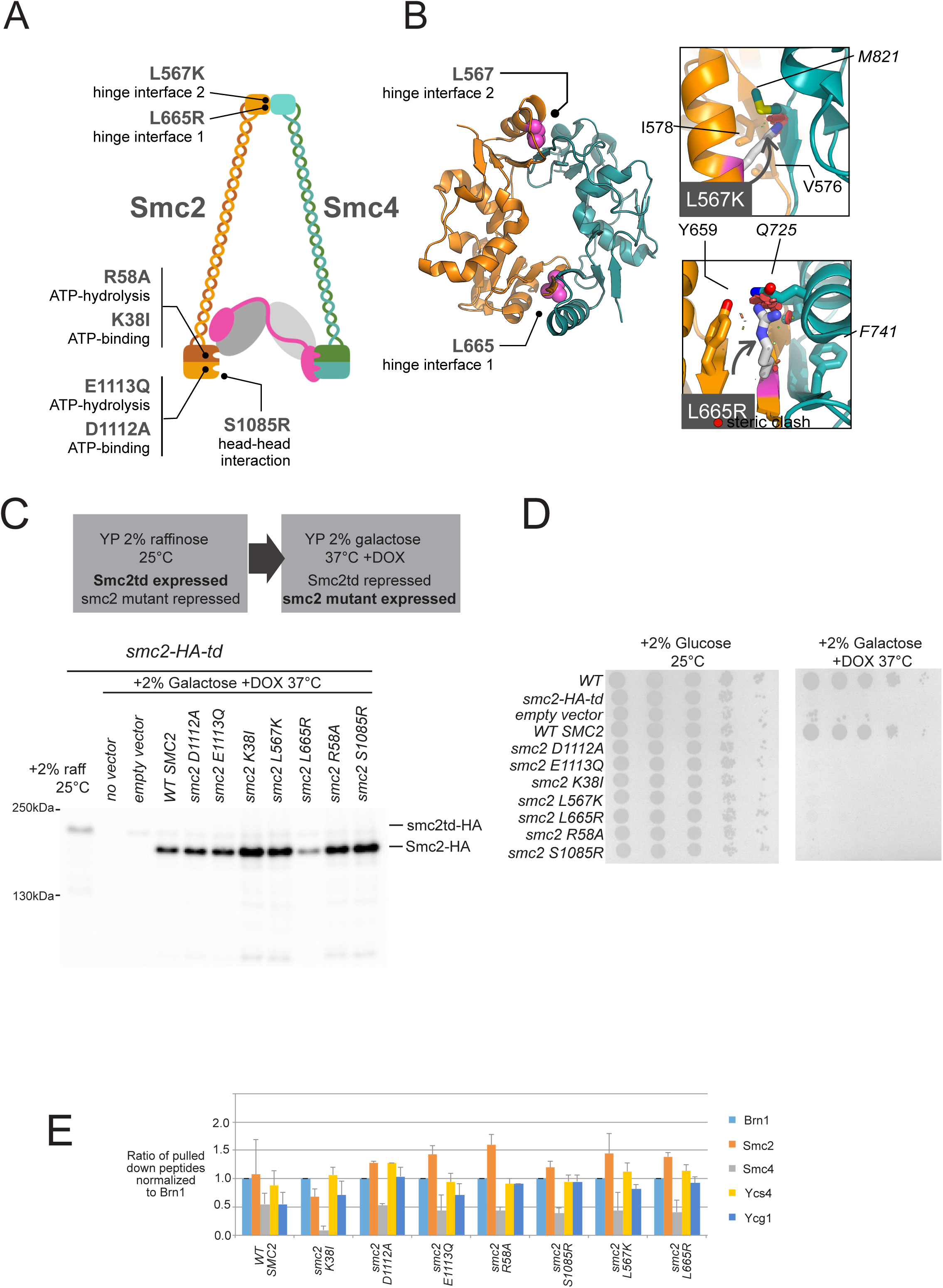
*Expression of inducible Smc2 mutants created by conditional protein exchange.* **A)** Schematic view of budding yeast condensin, illustrating the relative locations of each Smc2 mutation. Engagement of ATP-bound head domains triggers ATP hydrolysis, leading to their disengagement. The K38I and D1112A mutations are predicted to block binding of ATP to their respective pockets. The E113Q mutation is predicted to slow down the rate of ATP hydrolysis, while R58A is predicted to disrupt DNA dependent ATP hydrolysis. The S1085R mutation is predicted to disrupt head-head interactions. **B)** Within the hinge region of Smc2, we designed two mutations to specifically disrupt either of the two interfaces made with Smc4: hinge interface 1, L665R; hinge interface 2, L567K. Inset panels show additional details for how each single point mutant is predicted to generate a series of steric clashes (red circles) incompatible with hetero-dimerization of the hinge. Key amino acids of Smc2 and Smc4 are labelled in plain and italic text respectively. Figures were generated using MacPyMOL. **C)** Conditional protein exchange system for *smc2* mutations. Under the permissive conditions (2% raffinose, 25°C) functional degron and HA tagged protein is expressed. Under the restrictive conditions (+2% galactose, + doxycycline 37°C), the functional degron protein is degraded and the exogenous HA tagged protein is expressed from a *GAL1* promoter. **D)** Spot test of viability following conditional protein exchange for the different *smc2* mutants. **E)** Condensin complexes generated following expression of different *smc2* mutants. Following conditional protein exchange, *in vivo* condensin complexes were immuno-precipitated and the isolated complexes analyzed by MS/MS as described in the Methods. The signal from several peptides derived from each protein was quantitated by MS/MS and then normalized to the quantity of peptides from the Brn1 subunit. This provides a measure of the relative quantity of each subunit immuno-precipitated by Brn1-V5. All experiments are the average of two independent experiments and the standard deviation of the experiment shown by the error bar.

Using this system, only GAL1p-driven expression of wild-type SMC2 rescued growth after Smc2 depletion; none of the mutants restored viability (Figure 1D).Therefore, each of our *smc2* mutations inhibit essential functions of condensin. Functionally, loss of condensin activity results in chromosome mis-segregation (Lavoie *et al*, 2000). We therefore examined the effects of these mutations on chromosome duplication and segregation over the course of a single cell cycle. Cells were arrested in G1 using alpha factor pheromone, before degradation of endogenous Smc2 and expression of mutant proteins. FACS analysis of DNA content of each strain demonstrated that all mutants completed DNA replication, generating 2C cells (Figure 3A). During chromosome segregation (80–100 minutes post-release), all *smc2* mutants produced aneuploid cells (Figure 3A). In contrast, expression of *WT SMC2* led to the normal generation of 1C daughter cells (Figure 3A). We conclude that following expression of mutant Smc2 proteins, budding yeast cells displayed the canonical loss of condensin function phenotype.

Loss of condensin function in Smc2 mutants may simply be due to disruption of the intact pentameric condensin complex. To test whether the complex was still intact following expression of each mutant we tagged the endogenous Brn1 kleisin subunit with the V5 epitope and immuno-precipitated the complex from cells using anti-V5 antibodies. We resolved the immuno-precipitated complex by SDS PAGE and analyzed the resolved proteins by MS/MS. In all mutations, apart from *smc2K38I,* peptides from each condensin subunit could be identified at similar ratios (Figure 1E). In *smc2K38I* a reduced ratio of Smc4 peptides relative to Brn1 was observed compared to the other subunits, suggesting that the K38I mutation inhibits Smc4 binding to the complex. To confirm that Brn1 and smc2K38I proteins could still interact we performed pull downs of Brn1-V5 in *smc2K38I* cells and performed western blots to compare the extent of Smc2 protein in the immunoprecipitated in *smc2K38I* cells compared to other smc2 mutant cells (Supp. Fig 1). We observed that the smc2K38I protein was pulled down with Brn1 to a similar extent as in other *smc2* mutants (Supp. Fig 1). Therefore, our data suggests that the stable Smc2/Smc4 heterodimer is sensitive to loss of ATP binding to the Walker A Smc2 ATP binding pocket in comparison to disruption of the Walker B motif. In conclusion we find that all the *smc2* mutations, except *smc2K38I*, are incorporated into a complete condensin complex.

Next, we sought to examine how these different mutations altered chromatin association of condensin. In budding yeast, condensin is strongly enriched at centromeres and within the rDNA repeats (Cuylen *et al*, 2011; D’Ambrosio *et al*, 2008; Verzijlbergen *et al*, 2014) and the chromosome conformation of these regions is selectively structured in early mitosis by condensin action (Lavoie *et al*., 2002; Schalbetter *et al*., 2017). Using ChIP RT-PCR (<u>Ch</u>romatin Immuno-<u>P</u>recipitation and <u>R</u>eal <u>T</u>ime <u>PCR</u>) analysis we confirmed that V5-tagged Brn1 was enriched at both sites relative to a chromosomal arm region (ACT1) and that enrichment at both centromeres and rDNA was dependent on an intact condensin complex since depletion of Ycg1 led to loss of enrichment (Supp. Figure 2A). Condensin enrichment at centromeres and rDNA was also observed when we used a different V5-tagged condensin subunit for ChIP (Ycg1, Supp. Figure 2B). Having established the robustness of our detection procedures, we next attempted to ascertain how the different putative stages of the condensin enzymatic cycle alter its chromatin binding.

**Figure 2.**
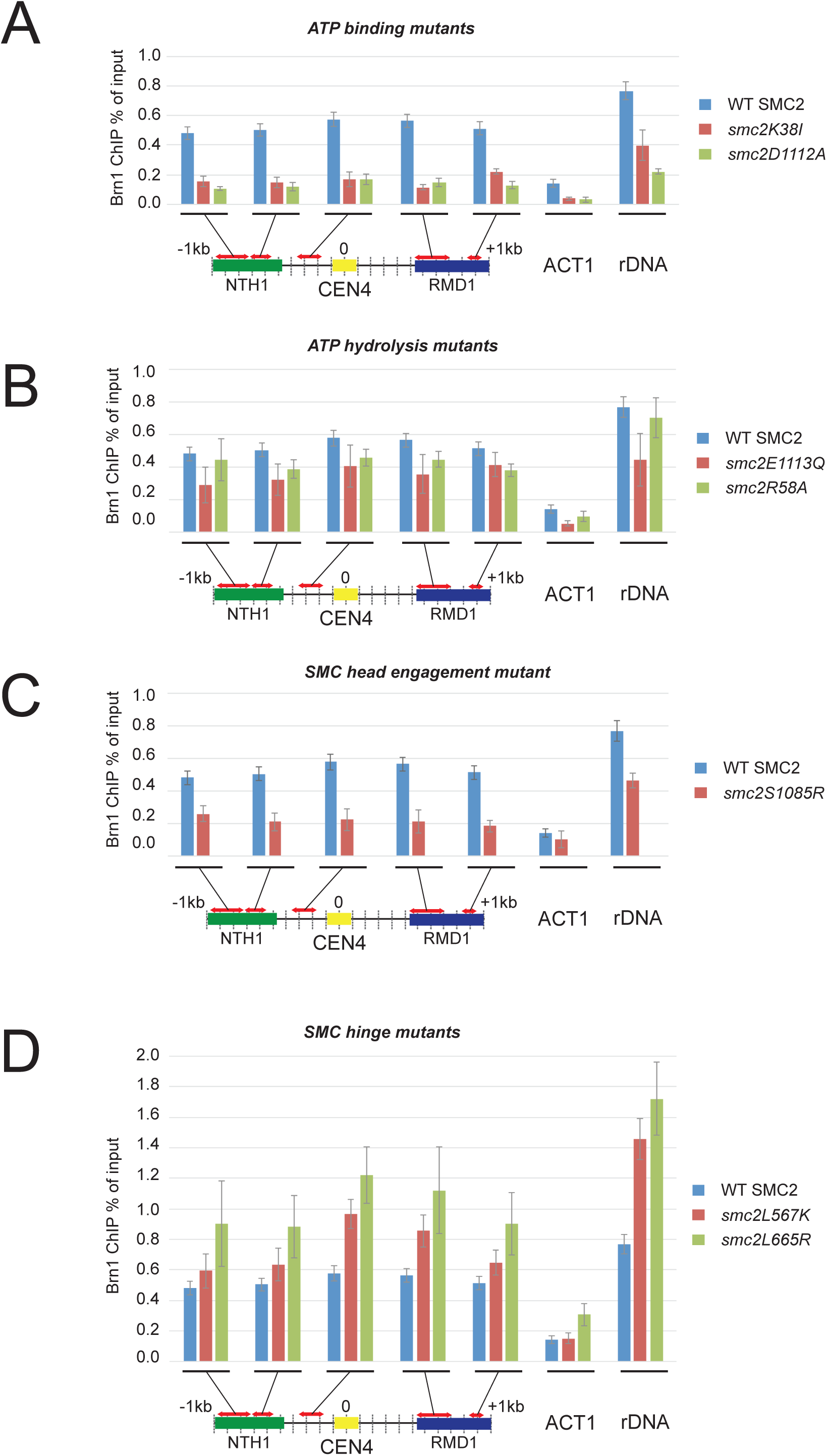
*Chromatin association of condensin following expression of smc2 enzymatic mutations.* Conditional expression depletion strains were arrested in G1 and endogenous Smc2 degraded and the indicated version of exogenous Smc2 expressed. Cells were then released from G1 into restrictive medium containing nocodazole. After 2 h, cells were harvested, and Brn1-V5 levels at the indicated sites analyzed by ChIP-qPCR. The average of at least three experimental repeats (qPCR performed in triplicate in each case) is shown with error bars. Enrichments are grouped according to function,

Expression of either the Walker A (*smc2k38I*) or the Walker B (*smc2D1112A*) ATP binding mutants led to loss of condensin ChIP enrichment across both the centromere and the rDNA (Figure 2A). Loss of condensin enrichment could also be observed at the *ACT1* locus indicating that even the low levels of condensin association with the chromosome arm were reduced following expression of these mutants. In contrast, expression of ATP-hydrolysis mutants led to a minor reduction in chromatin association compared to wildtype Smc2 (Figure 2B). We conclude that ATP binding but not ATP hydrolysis is required for stable association of condensin with chromatin. The destabilization of Smc4 binding to complex in the *smc2k38I* mutant would account for the loss of chromatin association (Figure 1E). However, we also found that the intact condensin complex formed in *smc2D1112A* cells does not stably associate with chromatin (Figure 2A). This indicates that ATP binding within an intact condensin complex is required for chromatin association. ATP binding is required for head-head engagement of BsSMC dimers (Hirano *et al*., 2001) whereas hydrolysis promotes head dis-engagement. The requirement for ATP binding in the intact condensin complex could be due to head-head engagement being a limiting step for stable chromatin association. To test this model, we generated a Smc2 mutation predicted to inhibit head-head engagement, S1085R (Figure 1A) (Hirano *et al*., 2001). Consistent with head-head engagement being required for chromatin association, expression of S1085R significantly reduced chromatin association (Figure 2C). Together, the cumulative evidence from both ATP binding, ATP hydrolysis and head engagement mutants, indicate that SMC ATP binding and head-head engagement are required for chromatin association of condensin.

We next investigated the role of the SMC-hinge in chromatin association. Models of hinge function indicate that a DNA binding surface at the hinge to coiled-coil transition point becomes exposed, following partial destabilization of the hinge interface (Alt *et al*., 2017; Hirano & Hirano, 2002, 2006; Soh *et al*, 2015). Expression of smc2L567K (disrupting interface 2) or smc2L665R (disrupting interface 1) increased condensin association at both centromeres and rDNA (Figure 2D). We conclude that a stable SMC hinge normally limits accumulation of condensin at centromeres and the rDNA.

The differential chromatin association of the different *smc2* mutations allowed us to assess if there was any connection between subunit phosphorylation and chromatin association. Previous studies have shown that mitotically phosphorylated Ycg1 has a retarded electrophoresis mobility (Lavoie *et al*., 2004; St-Pierre *et al*., 2009). We observed a mitotic specific mobility shift of Ycg1 in cells expressing wildtype Smc2 and the two hinge mutants (*smc2L567K* and *smc2L665R*) (Figure 3B). In this assay the extent of Ycg1 mobility-shift caused by ATP binding, ATP hydrolysis mutants, and head engagement mutants was more difficult to discern (Figure 3B). This analysis suggested that Ycg1 subunit phosphorylation is extensive under conditions when condensin is stably bound to chromosomes but diminished when binding is unstable. Next, we examined the number of distinct condensin phospho-peptides identified by mass spectrometric analysis of isolated condensin complexes, compiling a table of phosphorylated peptides identified with >95% certainty over the course of two independent experiments (Table 1). By plotting the number of unique phosphorylated peptides detected, against the extent of chromatin enrichment at CEN4 for each mutant (Supp. Fig. 3), we found a correlation between overall extent of phosphorylation of condensin and chromatin association (R^2^ = 0.83) (Supp. Fig. 3A). Looking at each subunit individually we also observed correlations between the extent of Ycg1 and Brn1 phosphorylation and chromatin association (R^2^ = 0.86 and R^2^ = 0.84) (Supp. Fig. 3B and C). For Ycs4 we observed a much weaker correlation (R^2^ = 0.47) (Supp. Fig. 3D). These data suggest that the extent of phosphorylation of Ycg1 and Brn1 scales with chromatin enrichment. To investigate this connection between condensin phosphorylation and chromatin binding further, we next investigated the importance of condensin subunit phosphorylation for chromatin association.

**Figure 3.**
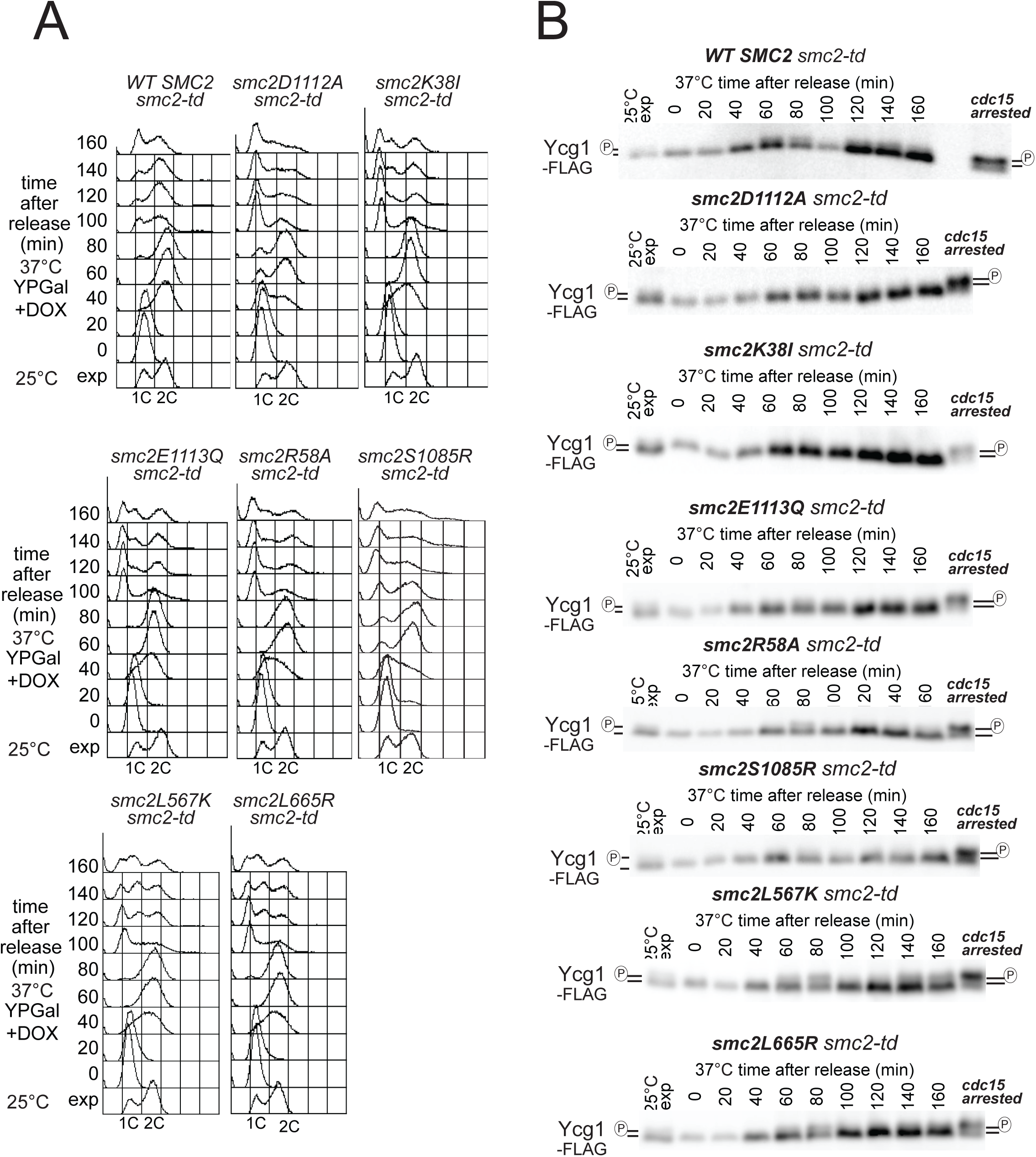
*Cell cycle profile and Ycg1 phosphorylation following expression of smc2 enzymatic mutations.* Conditional expression depletion strains were arrested in G1 and endogenous Smc2 degraded and the indicated version of exogenous Smc2 expressed. Cells were then released from G1 into restrictive medium containing nocodazole. Every 20 minutes samples were taken for **A)** FACS analysis of DNA content **B)** Western blotting for assaying the phospho-shift of FLAG-tagged Ycg1. Samples of wildtype FLAG-tagged Ycg1 cells arrested at late mitosis (*cdc15* arrested) is shown to demonstrate the extent of Ycg1 phospho-shift in late mitosis.

Sites of serine/threonine phosphorylation in condensin subunits have been extensively mapped in budding yeast. Smc4 contains all functional CDK phosphorylation sites in yeast condensin (Robellet *et al*., 2015). Brn1, Ycg1 and Ycs4 all contain Cdc5/Polo dependent phosphorylation sites (St-Pierre *et al*., 2009). Using previously characterized S/T to A mutants of mapped phosphorylation sites (Robellet *et al*., 2015; St-Pierre *et al*., 2009), we assessed how condensin phosphorylation regulated condensin chromatin association. In FRAP assays CDK mediated phosphorylation of budding yeast Smc4 stabilizes the association of the protein with chromatin (Robellet *et al*., 2015; Thadani *et al*., 2018). To confirm this finding, we assayed the condensin complex chromatin enrichment in yeast where the potential CDK sites of Smc4 had been mutated to alanine – *smc4-10A* (Robellet *et al*., 2015). Cells containing this mutation displayed loss of condensin enrichment at both the centromere and the rDNA (Figure 4A). Next, we examined strains where all the mapped phosphorylation sites on the non-SMC subunits had been mutated to alanine, specifically yeast strains *ycg1-521*, *brn1-570* and *ycs4-543*. Condensin enrichment at both centromeres and the rDNA was significantly reduced in cells containing either the *ycg1-521* or the *brn1-570* alleles (Figure 4B and C). Enrichment was only partially reduced in *ycs4-543* (Figure 4D), consistent with the poorer correlation between extent of phosphorylation and chromatin association (Supp. Fig. 3D). This data supports a model where condensin phosphorylation in budding yeast generally facilitates chromatin association and accumulation of chromatin bound condensin. Next, we sought to examine the kinases required for chromatin enrichment. In several contexts, Aurora kinase is required for chromatin association of condensin (Giet & Glover, 2001; Hagstrom *et al*, 2002; Petersen & Hagan, 2003). However, Ipl1 function is not required for the chromatin enrichment of condensin in late mitosis (Lavoie et al., 2004). Notably, loss of function of the Aurora kinase Ipl1 delays the mitotic phosphorylation dependent mobility shift of the non-SMC condensin subunits Brn1, Ycg1 and Ycs4, but does not prevent their extensive phosphorylation in late stages of mitosis (St-Pierre *et al*., 2009). We therefore examined whether loss of Ipl1 altered condensin chromatin association in early mitosis, when phosphorylation of condensin is relatively low in *ipl1* cells. Loss of Ipl1 resulted in significant loss of condensin enrichment across the centromere but only a relatively minor change at the rDNA (Figure 5A). Therefore, in early mitosis Ipl1 activity is required for condensin enrichment at centromeres, consistent with the cellular localization of Ipl1 at this stage of mitosis (Biggins & Murray, 2001). Both *in vivo and in vitro* analysis has indicated that, while Ipl1 regulates non-SMC phosphorylation in early mitosis, Cdc5/Polo is generally required for extensive phosphorylation of non-SMC condensin subunits (St-Pierre *et al*., 2009).To test if Cdc5 activity was required for Ipl1 dependent chromatin association we assayed condensin enrichment following Cdc5 inactivation through two distinct *ts* alleles, *cdc5-10* and *cdc5-99* (Figure 5B, C). Surprisingly, we did not observe any significant loss of condensin enrichment at either the centromere or rDNA in either *cdc5* mutant strain at this early stage of mitosis. Therefore. the association of condensin at centromeres in early yeast mitosis is dependent on Ipl1/Aurora B but independent of Cdc5/Polo.

**Figure 4.**
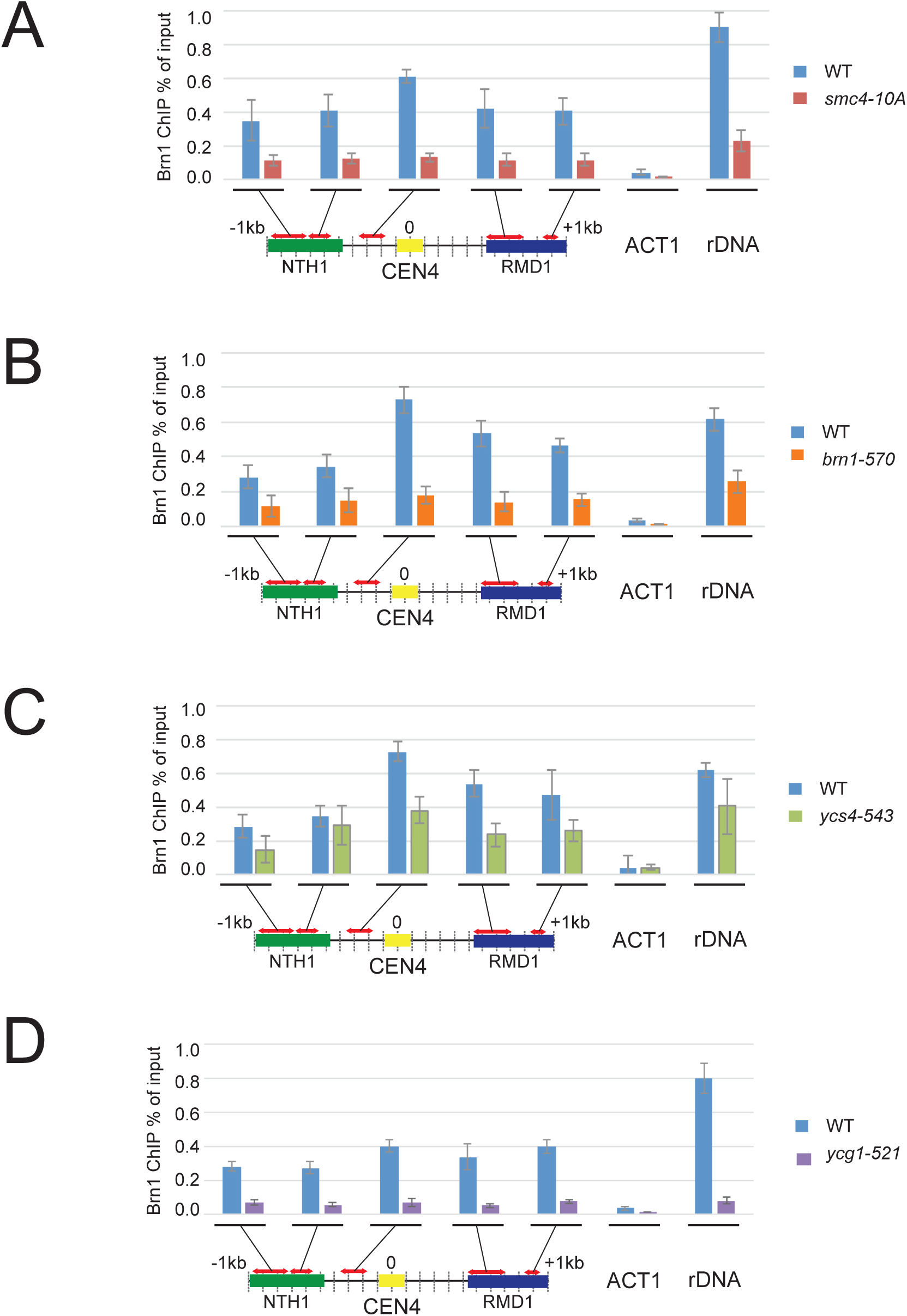
*Effects of phosphorylation defective condensin subunits on chromatin association.* Strains expressing the defined phosphorylation mutants of **A)** Smc4-*smc4-10A*, **B)** Brn1-*brn1-570*, **C)** Ycs4 – *ycs4-543* **D)** Ycg1, *ycg1-521* were released from G1 into medium containing nocodazole at 37°C. After 2 h, cells were harvested, and Brn1-V5 levels at the indicated sites analyzed by ChIP-qPCR. The average of at least three experimental repeats (qPCR performed in triplicate in each case) is shown with error bars.

**Figure 5.**
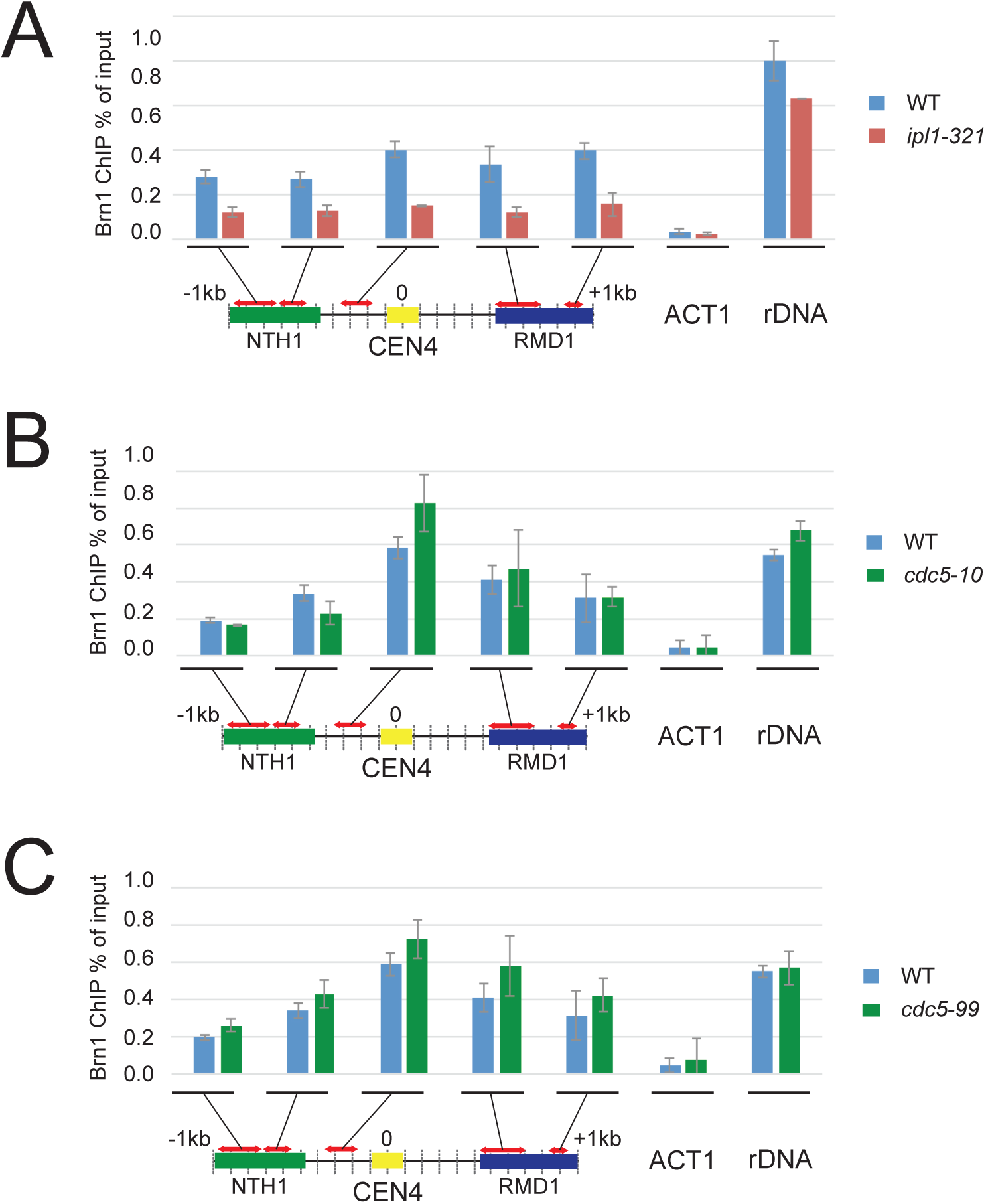
*Requirement for Ipl1 and Cdc5 for condensin chromatin association and complex stability.* Either **A)** *ipl1-321*, **B)** *cdc5-10* **C)** *cdc5-99* cells were released from G1 into medium containing nocodazole at 37°C. After 2 h, cells were harvested, and Brn1-V5 levels at the indicated sites analyzed by ChIP-qPCR. The average of at least three experimental repeats (qPCR performed in triplicate in each case) is shown with error bars.

## Discussion

Here we investigate the importance of both condensin enzymatic action and condensin protein phosphorylation for condensin chromatin association. Our results indicate that ATP binding and head engagement regulate budding yeast condensin’s chromatin binding *in vivo*. Conversely, we observe that stable hinge-hinge interactions limit condensin accumulation at centromeres and rDNA.

ATP binding appears to promote chromatin association through two mechanisms: stabilizing Smc4 binding to the complex (lost in *smc2K38I*) and enabling ATP-dependent head engagement. Head-head engagement is proposed to be a key step in DNA association of all SMCs (Burmann & Lowe, 2023). Our observation that head-head engagement is required for chromatin association aligns budding yeast condensin with previous studies showing that BsSMC mutations that disrupt head engagement also inhibit DNA binding both *in vitro* and *in vivo* (Hirano *et al*., 2001; Minnen *et al*., 2016). However, we should note that another study examining head-head engagement mutations in in budding yeast condensin did not observe a decrease in chromatin engagement of condensin (Thadani *et al*., 2018). This discrepancy may reflect differences in culture temperature in the two studies, with the negative effect of head engagement mutations evident at 37°C but not at 30°C.

Our data also show that chromatin association is only the first step of condensin action on chromosomes. Other condensin mutants, unable to efficiently hydrolyze ATP or maintain a stable hinge, still bind to chromatin but create an inactive state. Disrupting the hinge engagement of Smc2 increased chromatin enrichment of the complex at the CEN and rDNA. These findings are consistent with previous studies that concluded that SMC hinge-hinge interaction is an important regulator of DNA binding and SMC activity (Alt *et al*., 2017; Hirano & Hirano, 2002, 2006; Soh *et al*., 2015). Models of how conformational changes at both the head and hinge of *Bs*SMC complexes regulate SMC complex activity have been postulated (Hirano & Hirano, 2006; Soh *et al*., 2015). DNA binding at the hinge could stimulate ATP hydrolysis events that take place following initial DNA interaction (Hirano & Hirano, 2006; Soh *et al*., 2015). One possible interpretation of our results is that partial disruption of the hinge allows an initial state of DNA interaction stabilized by head engagement but inhibits movement of the initially loaded complex from its loading site. This would prevent movement of condensin away from the loading sites at the centromere or the RFB and cause heightened accumulation of condensin at these loci. Such a model would be analogous to accumulation of cohesin at the centromere when its enzymatic cycle is inhibited down stream of ATP binding (Hu *et al*, 2011; Petela *et al*, 2018; Srinivasan *et al*, 2018). Whatever the exact mode of action, our data indicate that enzymatic control of SMC-DNA interactions through head-head and hinge-hinge interactions is conserved across bacterial and eukaryotic SMCs.

We also find that condensin phosphorylation correlates with enzymatic state and chromatin enrichment, suggesting it promotes chromatin association. Others have suggested that condensin subunit phosphorylation provides a mechanism to relieve the repression of condensin enzymatic cycle caused by unstructured regions in the vicinity of the SMC head domains of the complex (Cutts *et al*., 2025; Tane *et al*., 2022). Such a scenario is consistent with the reduced levels of chromatin association we observe following expression of Ycg1 and Brn1 phosphorylation deficient proteins. The correlation we observe between chromatin association of the condensin complex and extent of phosphorylation of Ycg1 and Brn1 is also consistent with the centromeric localization of the condensin regulators Ipl1 and Cdc5 (Garcia-Rodriguez *et al*, 2019; Mishra *et al*, 2016). Consistent with the close connection of chromatin association and phosphorylation, we find that active Aurora/Ipl1 kinase promotes condensin phosphorylation and chromatin association at centromeres in early mitosis. In fission yeast Aurora kinase directly targets condensin to facilitate chromatin binding (Tada *et al*., 2011). Surprisingly, we find that Cdc5 activity is not required for chromatin association, at least at centromeres and rDNA in early mitosis. Cdc5 is recruited to chromatin by cohesin complexes and is required for rDNA chromosome compaction in mid to late mitosis (Lamothe *et al*, 2020). At centromeres, phosphorylation of Polo kinase sites could be redundant with other, kinetochore localized, kinases, for example Mps1 (Kagami *et al*, 2014) or Ipl1 may target condensin in a manner that promotes chromatin association independently of Cdc5 early in mitosis. In this scenario Cdc5 phosphorylation of condensin in budding yeast could promote condensin loading later in mitosis. Alternatively, phosphorylation may act both upstream and downstream of chromatin loading, facilitating turnover of the enzymatic cycle. Here, Cdc5 may act downstream of loading, potentially to promote turnover of partially phosphorylated complexes. Constitutive phosphorylation of budding yeast condensin CDK sites reduces both turnover on chromatin and compromises rDNA mitotic compaction (Thadani *et al*., 2018) The hyper-phosphorylation we observe on chromatin bound, hinge destabilized, condensin complexes could indicate accumulation of phosphorylation on chromatin complexes that are inhibited from progressing to the next stage of the enzymatic cycle by hinge destabilization.

In summary, our data indicate that yeast condensin’s enzymatic cycle regulates chromatin association similarly to other SMC complexes across prokaryotes and eukaryotes: ATP binding and head engagement initiate stable chromatin interaction, while hinge dynamics and phosphorylation modulate subsequent steps. We also find that condensin phosphorylation is closely linked to chromatin association and accumulates on chromatin bound condensin complexes.

## Acknowledgements

We thank Luis Aragon, Sue Biggins and Damien D’Amours for yeast strains. This work was funded by the Biotechnology and Biological Sciences Research Council United Kingdom, (BBSRC UK) Grant ref BB/J018554/1 (S.A.S., M. Y., C. M), an MRC Career Development Award G1100257 (S.S.) and the Royal Society U.K. (J. B.).

## Data Availability

The mass spectrometry proteomics data have been deposited to the ProteomeXchange Consortium (http://proteomecentral.proteomexchange.org) via the PRIDE partner repository with the dataset identifier PXD006028.

## Materials and Methods

### Generation of smc2 mutants

#### Plasmid construction

pFA6a-Gal1-2xHA-6xHA-TRP1 (Baxter and Diffley, 2008) has been modified as follows: a linker fragment has been cloned near the BamHI site using InFusion with primers InFusion_pFA6_linker_F(GGACTATGCAGGATCtGGTGCTGGATCTGGTGGATTAATTAAaGGATCCATGGCGCCGGAGCAGGTTCTGGAGGTGCCGGTGGAGGCTATCCTTATGA) and InFusion_pFA6_linker_R(CATATGGATAGGATCTCAGCCGGCATAATCTGGCACG TCGT). The N-terminal 2x HA tag has been removed by cutting with PacI and religating. To ensure that there is no residual N-terminal tag, the upstream ATG has been removed by site-directed mutagenesis using the primers pFA6_ATG_SDM_F (GGAGAAAAAACCCGGAtctcaaattgtctttaattaaaGGATCCATggcgccGGA) and (pFA6_ATG_SDM_R

TCCggcgccATGGATCCtttaattaaagacaatttgagaTCCGGGTTTTTTCTCC). The SMC2 mutants which have been synthesised by Gene Cust and cloned into the pUC57 BamHI site. The *wt* and mutated *SMC2* constructs were subcloned from pUC57 into the modified pFA6 plasmid using the BamHI and SfoI sites. The resulting integration plasmid has been integrated into the TRP1 locus by recombination induced by cutting the plasmid with XbaI and transforming it into the yeast *smc2-td* cells.

### Strains used

All strains used are listed in Supplemental Table 2

### Cell Culture

Cell cultures were grown to log phase at 25°C in YP 2% raffinose, then arrested in G1 by the addition of alpha factor peptide (GenScript custom synthesis) 10 μg/ml. Once 90% of cells were in G1, Galactose (gal) was added to the cell culture (2%) followed by 50μg/ml doxycycline 15 minutes later. 30 mins after the addition of galactose, cell cultures were incubated at 37°C for 1 hour then released from G1 into depletion media (YP2%Raf/2%Gal + 50 μg/ml Dox 37°C). Nocodazole was added (10 μg/ml) at 40 min. Once 80-90 % cells were arrested in G2/M, cells were fixed in 1.5 % formaldehyde for 15 minutes. To quench the crosslinking reaction, 140 μl/ml of 1M glycine was added for 5 minutes. Cells were then washed once in PBS before the pellet was frozen in liquid nitrogen and stored at −80°C. For conditional protein depletion of Ycg1 using the auxin dependent protein degradation system, 0.5mM NAA (1-naphthaleneacetic acid) was added 15 minutes after the first wash from alpha factor.

### Chromatin Immunoprecipitation

Fixed cells were defrosted and re-suspended in 100 μl ChIP buffer (150 mM NaCl, 50 mM Tris HCl, 5mM EDTA, 0.5 % NP-40 (IGEPAL), 7 % Triton X-100, cOmplete Tablets, Mini EDTA-free EASYpack (Roche). Cells were lysed in a FASTPREP machine, 6 rounds of 30 seconds at 6.5 power, with 1 ml of 0.5 mm silica beads. The lysate was isolated by centrifugation and resuspended in 1ml ChIP buffer, before sonication for 15x 30 seconds (Biorupter Pico, Diagenode). 100 μl of sonicate was processed as an input control while 200 μl was incubated with 12.5μg/ml anti-V5 antibody (MCA1360, abD Serotec). For immuno-precipitation (IP) tubes were agitated for an hour and a half at 4°C with 45μl magnetic beads, (Dynabeads protein G – Life Technologies), washed 3 times in 1ml ChIP buffer were added and tubes incubated at 4°C for 2 hours.

Magnetic beads were isolated and washed four times in ChIP buffer, and a fifth time in ChIP buffer minus protease-inhibiting supplement. To reverse cross-linking, magnetic beads were incubated with 10% Chelex100 resin beads (BioRad 142-1253), in purified water at 95°C for 30 min. Samples were spun down and the supernatant kept at −20°C prior to analysis by qPCR. Input controls were precipitated using 0.1 x vol 3M NaAC pH5.2 and 2.5 x vol 100% EtOH and then cross-linking reversed as above, before purification with Nucleospin PCR clean up kit and eluted in nuclease-free purified water.

The immuno-precipitated DNA was analyzed using 2X AB-1323/B ABsolute™ QPCR SYBR. Green Low ROX Mix and processed in an MX3005p qPCR machine. Data was analyzed using the ‘Percentage Input Method’ where the CT values obtained from the ChIP are divided by the CT values obtained from the Input control samples. Since 1% of starting chromatin was used for each input sample, in order to adjust the CT value of input samples to 100% 6.644 (log2 of 100) was subtracted from it. Then the following formula was used to calculate the percentage input for each IP sample: 2^(adjusted ChIP input CT value – IP CT value) x 100. The percentage of input values obtained from the qPCR analysis of a minimum of three separate ChIP experiments were then averaged. Wildtype experiments performed alongside test were used for comparison. Error bars used for ChIP qPCR histograms represent the standard error of the mean.

### Condensin Immunoprecipitation

Immunoprecipitation (IP) buffer contained 50mM HEPES pH 7.5, 300mM KCL, 5mM EDTA, 0.5 % NP-40 (IGEPAL), 10% Glycerol, 80mM b-glycerophosphate, 0.01% Triton X-100, cOmplete Tablets, Mini EDTA-free EASYpack (Roche).

Cells were lysed in a FASTPREP machine, 5 rounds of 60 seconds at 6.5 power, with1 ml of 0.5 mm silica beads. Lysate spun out and the supernatant incubated with 12.5μg/ml anti-V5 antibody (MCA1360, abD Serotec) for 3-4 hours, rotating at 4°C. 50μl magnetic beads added, (Dynabeads protein G – Life Technologies), and left overnight rotating at 4°C.

Magnetic beads were isolated and washed three times in IP buffer, followed by resuspension in 1x SDS sample buffer ready for running on a SDS gel.

### Sample Preparation for Mass Spectrometry

Samples were run on SDS gels (4–15% Mini-PROTEAN® TGX™ Precast gel, Bio-Rad). Following resolution gels were Coomassie stained. Bands were excised, destained (50%MeCN, 25mM NH_4_HCO_3_) and dehydrated (speedvac). Followed by rehydration (10 mM DTT, 25mM NH_4_HCO_3_) and alkylation (55 mM chloroacetamide, 25mM NH_4_HCO_3_). Then destained and dehydrated as before, followed by an overnight in-gel trypsin digestion (12.5 ng/ µl; Sequencing grade modified Trypsin; Promega). Digested samples were then acidified by addition of trifluoroacetic acid (0.5% final volume).

### Mass spectrometry and data analysis

Peptide samples were analyzed by nano-LC-MS (ThermoFisher U3000 nanoLC and Orbitrap XL mass spectrometer). Peptides were loaded onto a C18 trapping cartridge (Pepmap100 C18; 0.3 x 5 mm i.d.; 5 μm particle size) for 5 min at a flowrate of 5 μL/min in 0.1% TFA loading buffer. Peptides were separated on an analytical column (PepMap100; 25 cm x 75 μm; 5 μm particle size) by a gradient from 1 to 35% acetonitrile over 28 min, in the presence of 0.1% FA, at a flow rate of 0.3 μL/min. Nanospray was from a New Objective emitter with 10 μm tip (FS360-20-10-N-20). Raw files were converted to .mgf format using MSConvert (Chambers et al., 2012) and a database search carried out using Mascot. The SwissProt database (Jan 2017) was employed, with species *S. cerevisiae* selected. Search parameters: 5 ppm precursor tolerance; 0.7 Da MS/MS tolerance; trypsin with up to 1 missed cleavage. Variable modifications: acetyl (protein N-terminus); deamidation (NQ); oxidation (M); phosphorylation (ST). Fixed modification: carbamidomethyl (C). Mascot peptide identifications were filtered to require an expectation value <0.05 and retain only top-ranking peptides.

Condensin subunit quantification was from MS1 (precursor) label-free quantification of 4-7 proteotypic peptides per subunit (Table 3) using extracted ion chromatograms in Skyline software (v3.6.1) (Schilling et al., 2012). Peptide intensities were summed from all gel slices for each mutant, averaged and normalized to Brn1 (epitope-tagged and used for pull-down).

### Western blotting and antibodies

Anti-V5 antibody (mouse monoclonal MCA1360, abD Serotec) used at 1:1000 Concentration in Western.

Anti-HA antibody (12Ca5 mouse monoclonal IgG_2_βK. Roche, Fisher scientific 10026563) used at 1:1000 in Western.

Anti-FLAG antibody (M2) SigmaAldrich.

## Supplemental information for Miles et al

**Supplemental Table 1.**
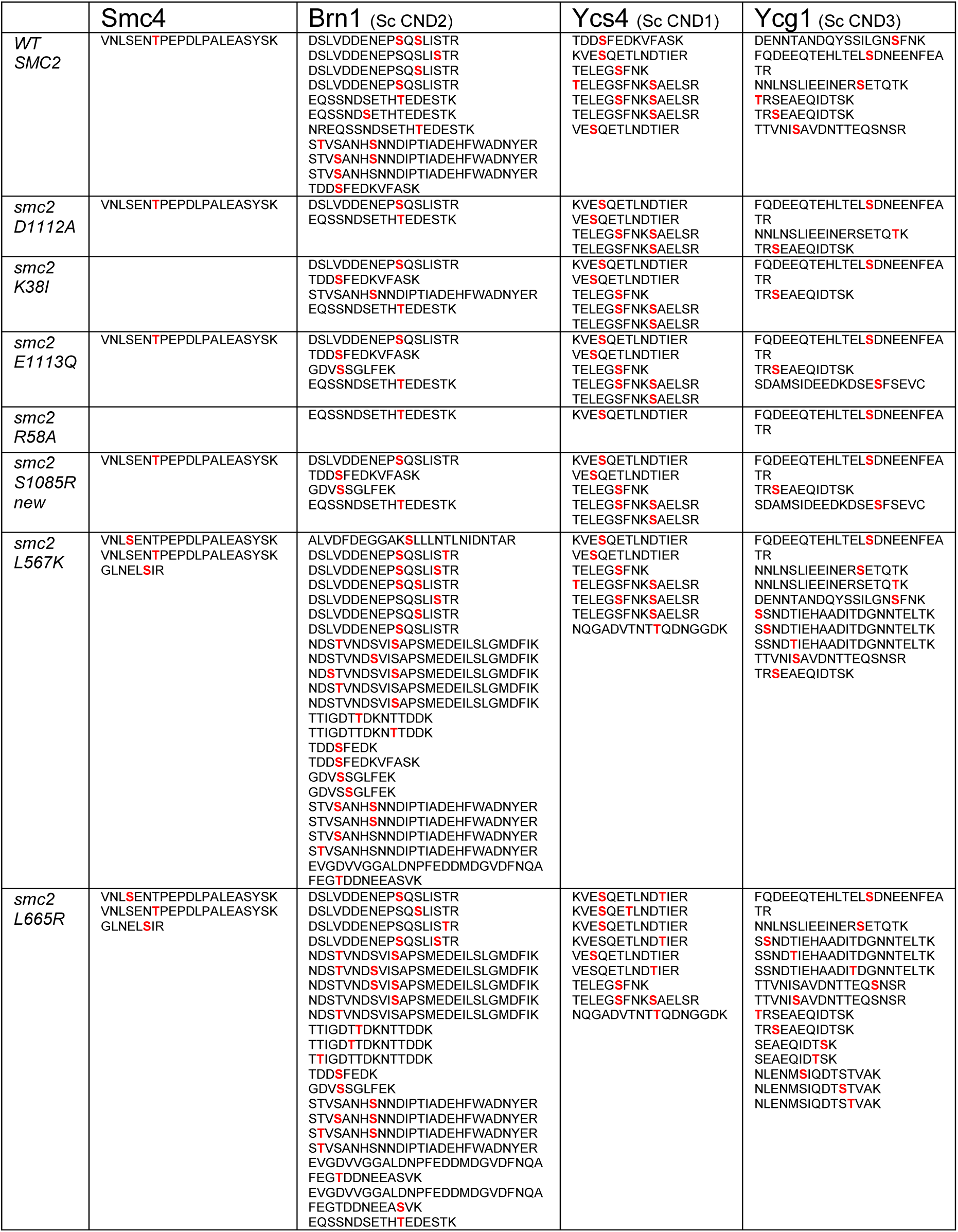
– Distinct phospho-petides identified following MS analysis from Brn1 immuno-precipitated complexes.

**Supplemental Table 2.**
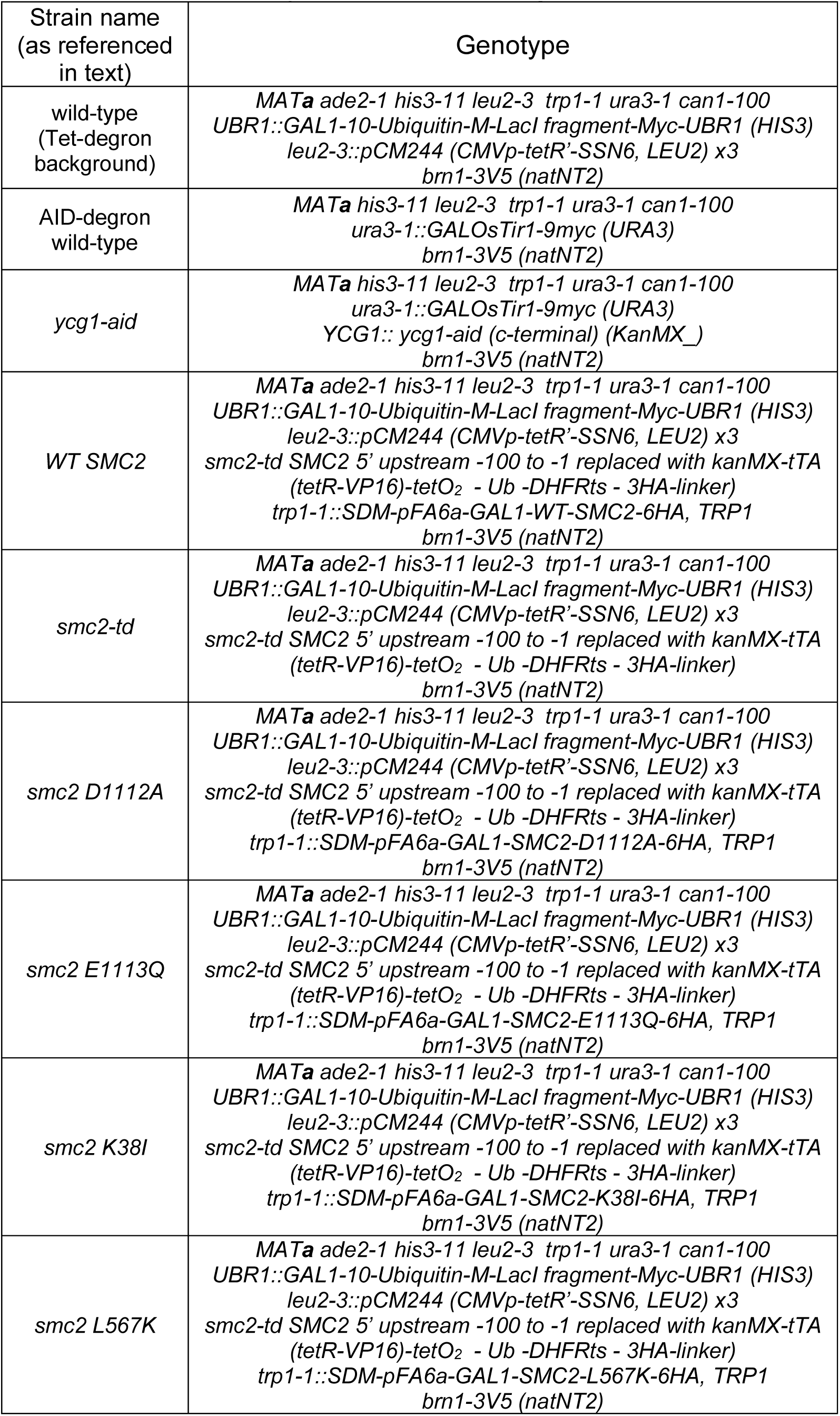
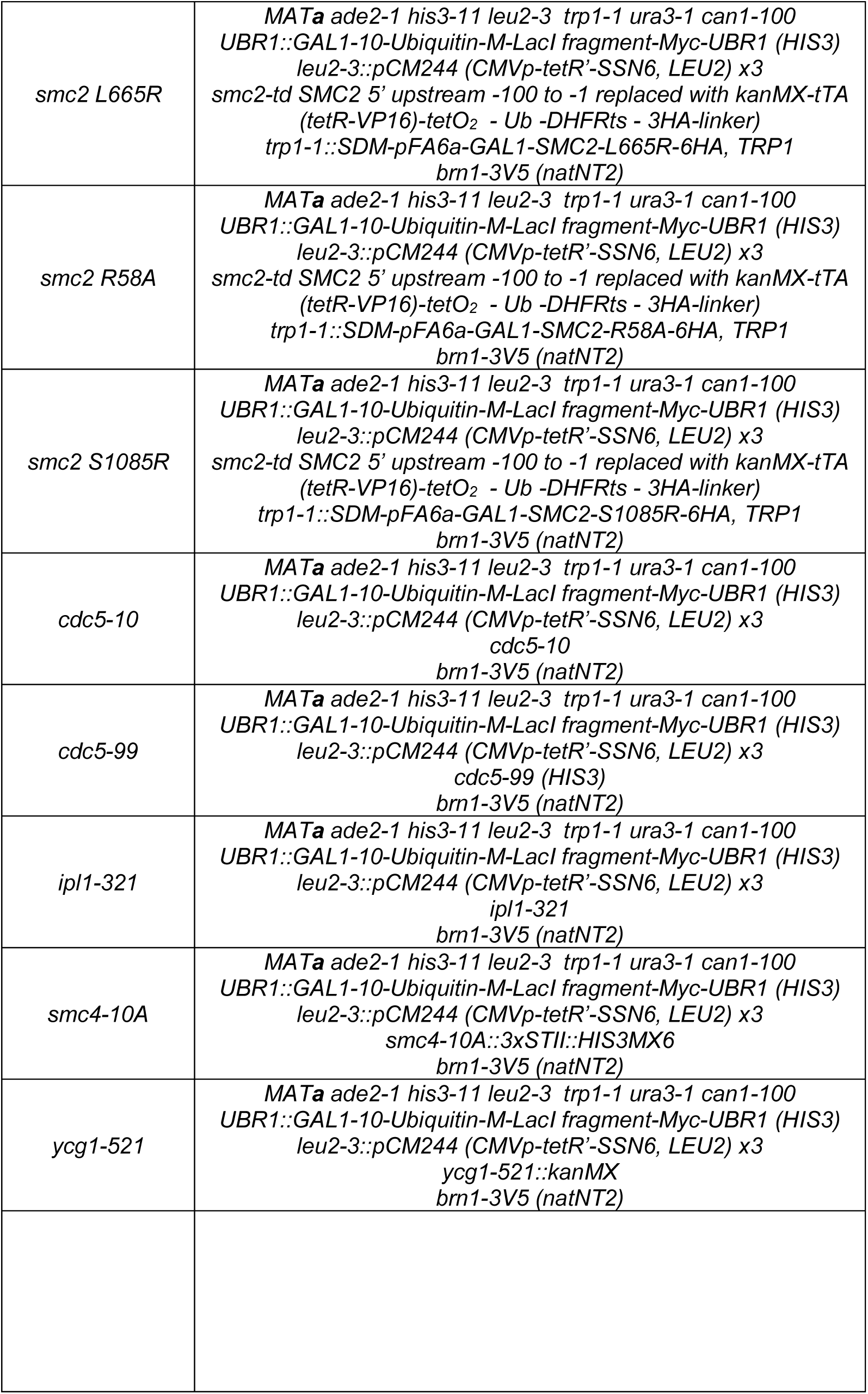
– Yeast strains used. All strains used in this study were of W303-1 background.

**Supplemental Table 3.**
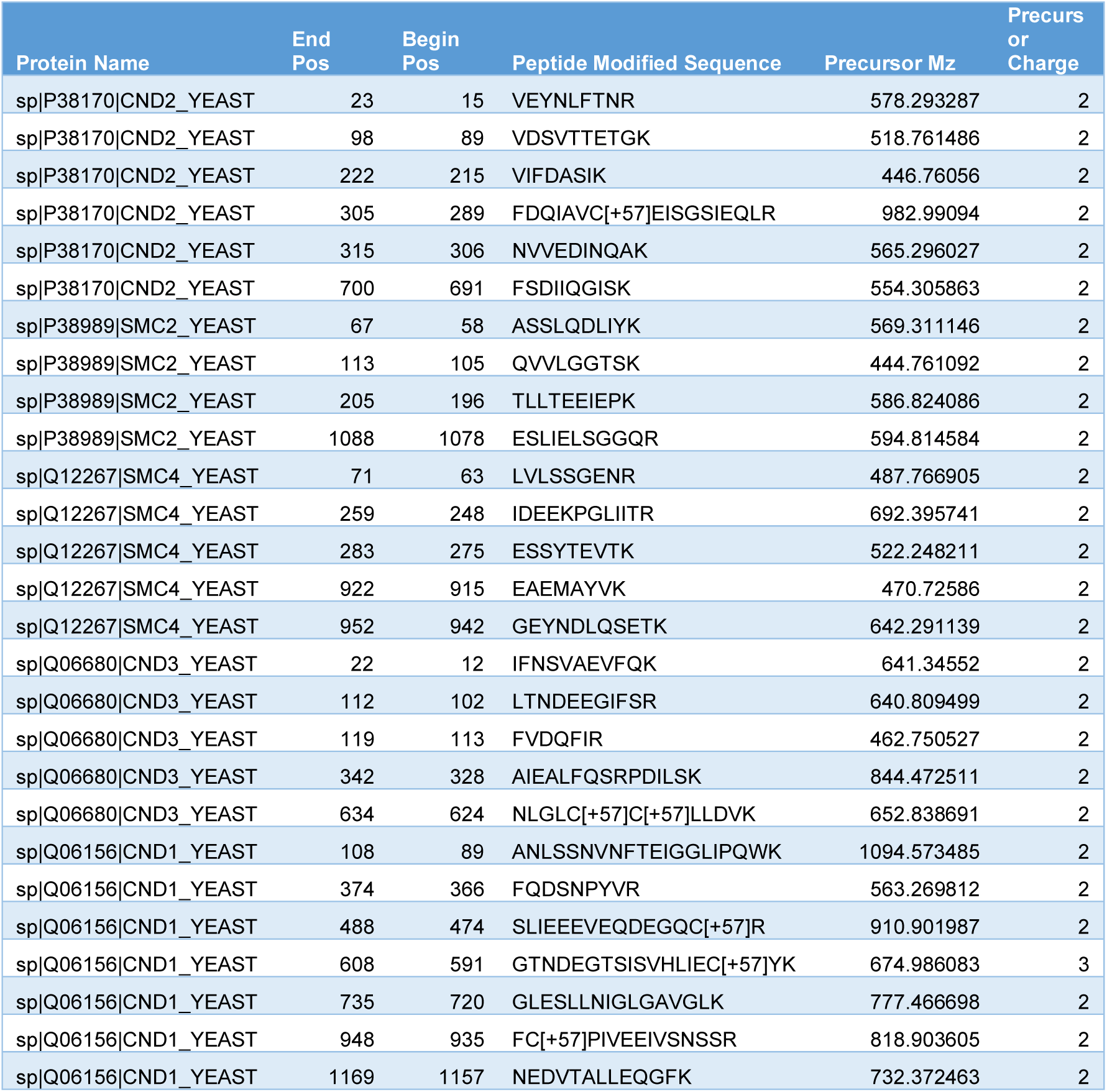
– Peptides used for label free quantification.

**Supplemental Table 4.**
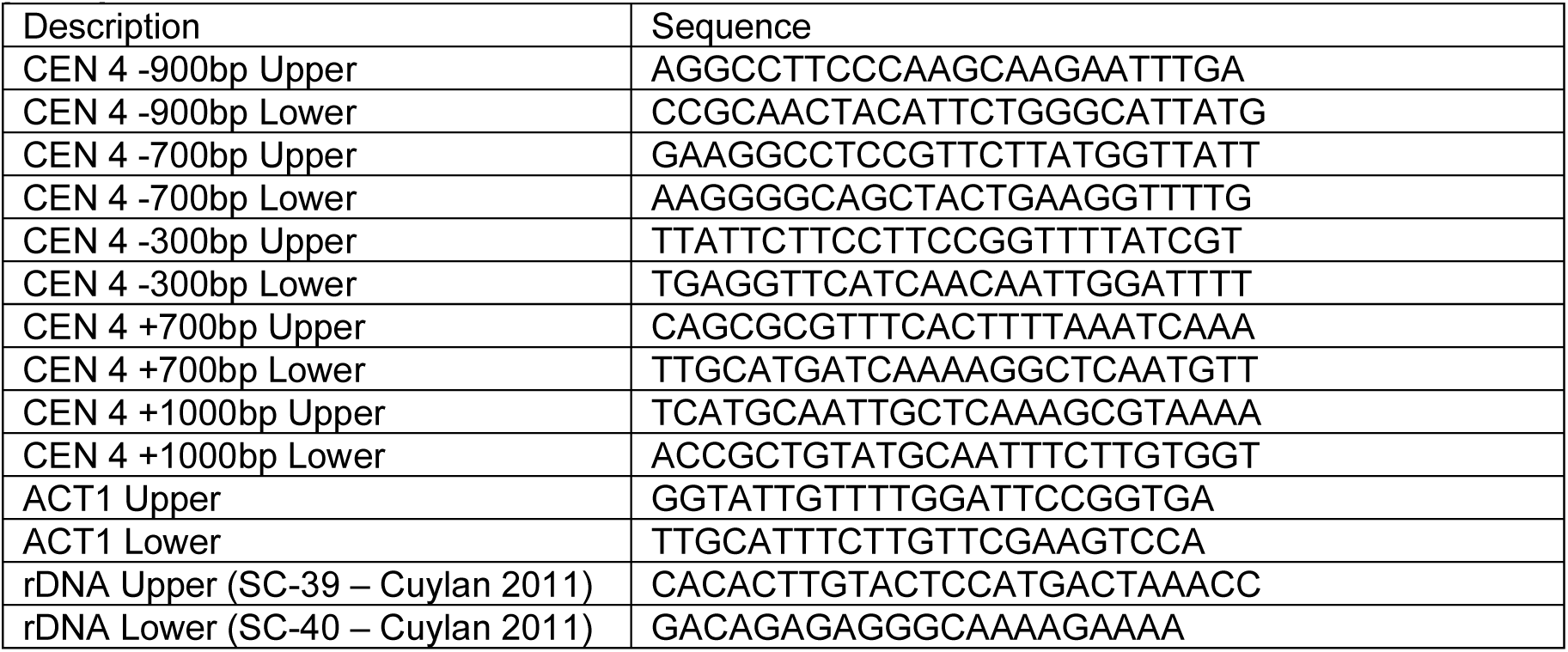
- Primers used for Real Time - PCR of immuno-precipitated chromatin.

**Supplemental Figure 1.**
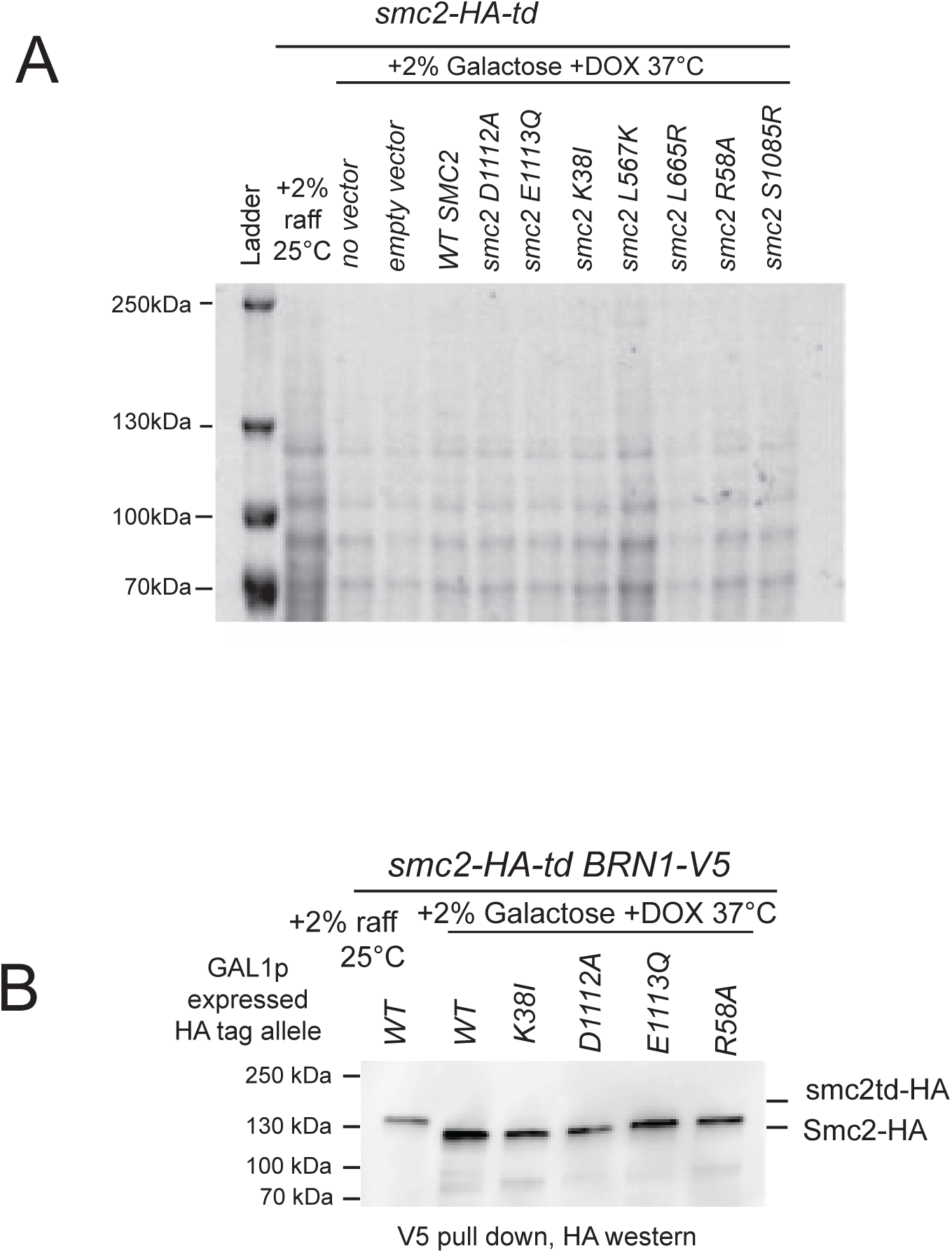
*Expression and immuno-precipitation of Smc2 proteins in smc2 enzymat-ic mutants* A) Image of the ponceau stained blot used in Figure 1C to show protein loading in lanes B) Equal numbers of cells were arrested in nocodazole following conditional protein exchange. Cell extracts were prepared and Brn1-V5 containing complexes immuno-precipitated and Western blotted for HA tagged Smc2.

**Supplemental Figure 2.**
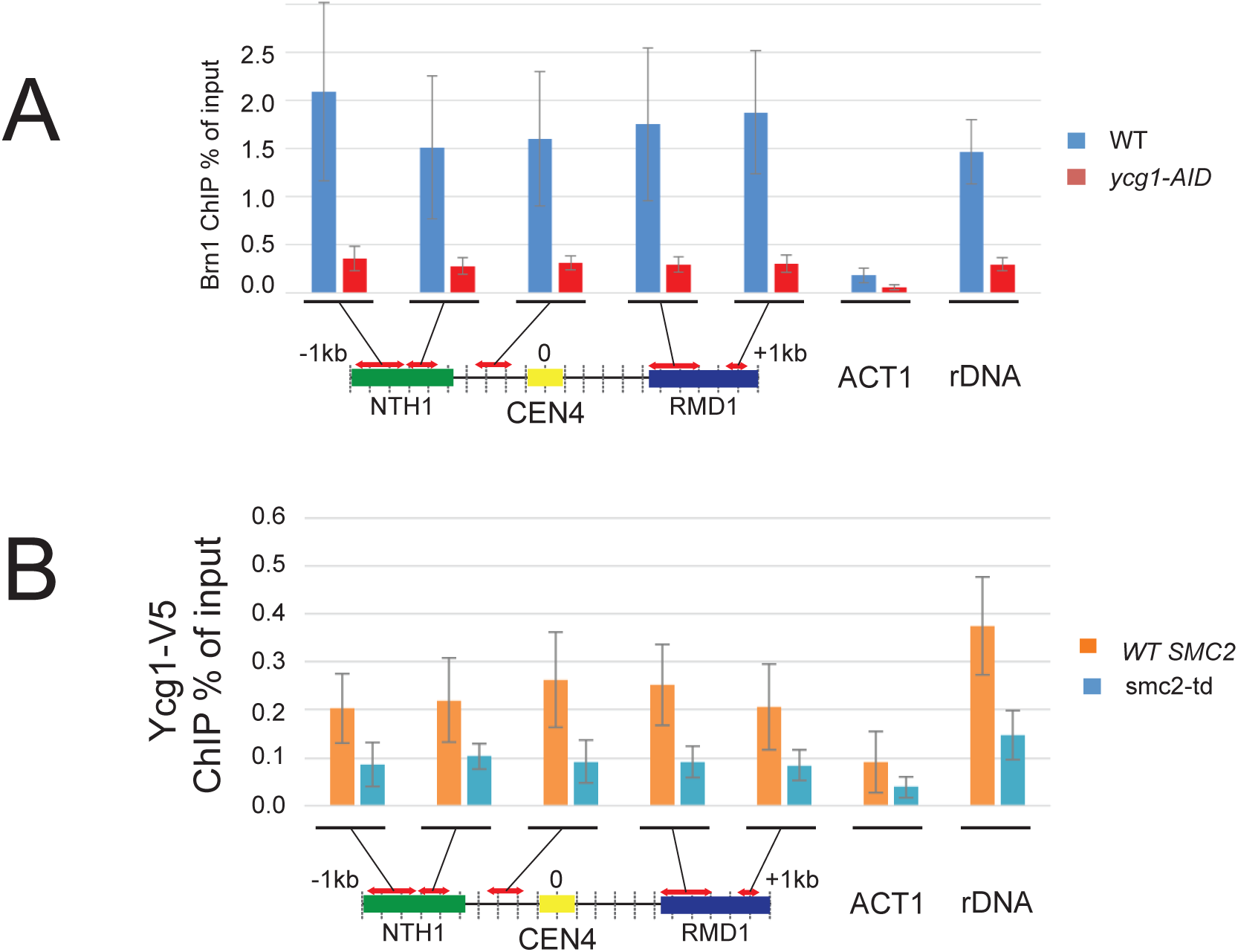
*Optimization of Condensin Chromatin Immuno-Precipitation at centromeres and rDNA in budding yeast* A) Histogram showing levels of Brn1-3V5 enrichment at specified loci surrounding CEN4 at the actin gene and the rDNA in either wildtype cells or cells depleted of Ycg1. Each sample was normalised to its corresponding input signal. Error bars represent SEM from three separate ChIP experiments. B) Histogram showing levels of Ycg1-3V5 enrichment at specified loci surrounding CEN4 at the actin gene and the rDNA in either GAL1-WTSMC2, smc2-td or smc2-td cells following galactose addition and degradation of smc2td protein. Each sample was normalised to its corresponding input signal. Error bars represent SEM from three separate ChIP experiments.

**Supplemental Figure 3.**
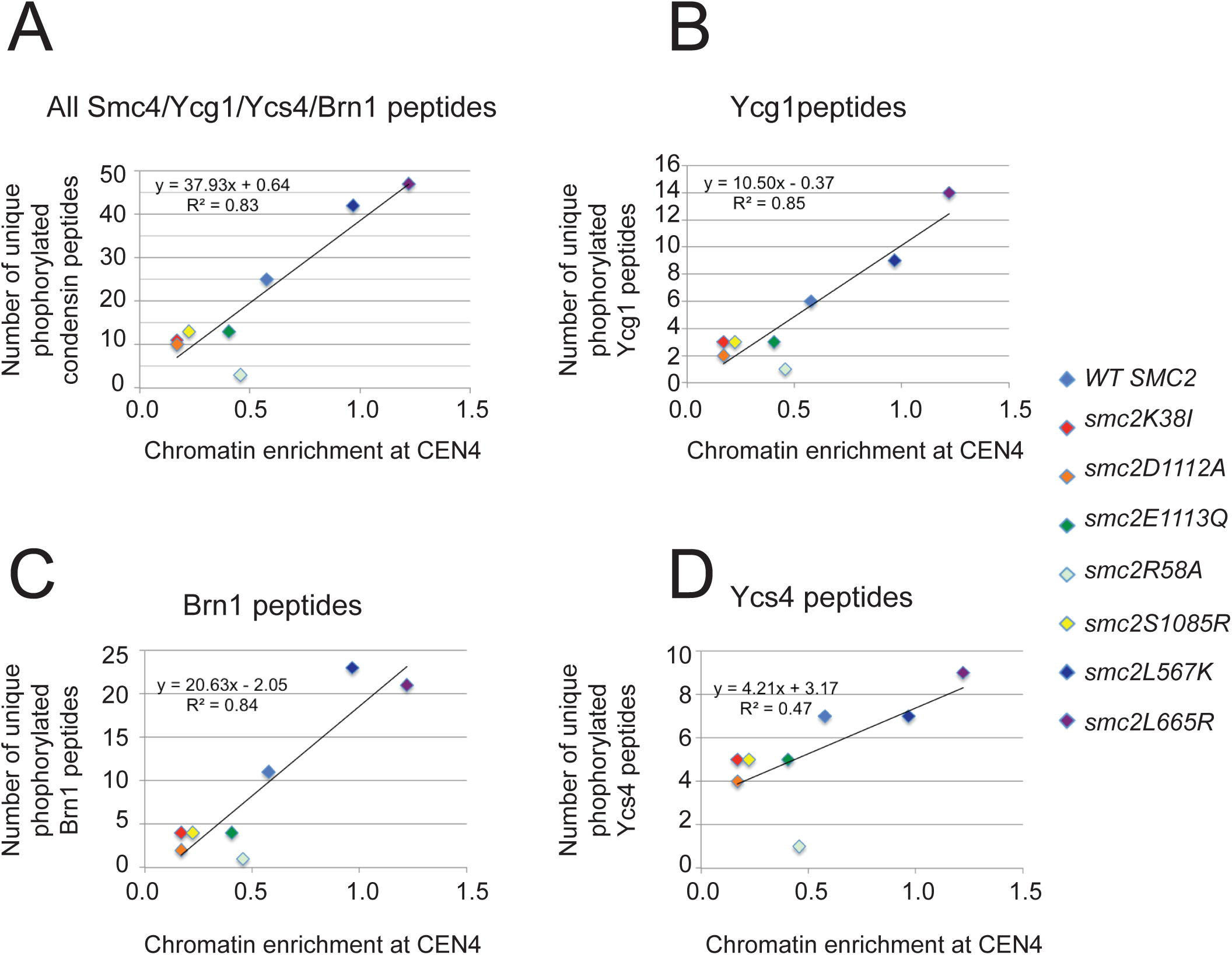
*Correlation between extent of phosphorylation of condensin and chroma-tin enrichment* Scattergrams are shown relating the number of distinct phosphorylated peptides identified from condensin subunits in each of the inducible SMC2 mutant strains (as listed in Supplementary Table 1) to the extent of chromatin enrichment at the Primer closest to centromere 4. Scattergrams are shown for peptides from all phosphorylated subunits **A)** Smc4/ Ycg1/Ycs4 and Brn1, **B)** For Ycg1 peptides, **C)** for Brn1 peptides **D)** for Ycg1 peptides. R^2^ for the best-fit line of each scattergram are shown. The key shows which SMC2 mutant each data point was derived from.

## References

1. Akai Y, Kurokawa Y, Nakazawa N, Tonami-Murakami Y, Suzuki Y, Yoshimura SH, Iwasaki H, Shiroiwa Y, Nakamura T, Shibata E et al (2011) Opposing role of condensin hinge against replication protein A in mitosis and interphase through promoting DNA annealing. Open Biol 1: 110023

2. Alt A, Dang HQ, Wells OS, Polo LM, Smith MA, McGregor GA, Welte T, Lehmann AR, Pearl LH, Murray JM et al (2017) Specialized interfaces of Smc5/6 control hinge stability and DNA association. Nat Commun 8: 14011

3. Aragon L (2018) The Smc5/6 Complex: New and Old Functions of the Enigmatic Long-Distance Relative. Annu Rev Genet 52: 89–107

4. Arumugam P, Gruber S, Tanaka K, Haering CH, Mechtler K, Nasmyth K (2003) ATP hydrolysis is required for cohesin’s association with chromosomes. Curr Biol 13: 1941–1953

5. Biggins S, Murray AW (2001) The budding yeast protein kinase Ipl1/Aurora allows the absence of tension to activate the spindle checkpoint. Genes Dev 15: 3118–3129

6. Burmann F, Clifton B, Koekemoer S, Wilkinson OJ, Kimanius D, Dillingham MS, Lowe J (2025) Mechanism of DNA capture by the MukBEF SMC complex and its inhibition by a viral DNA mimic. Cell 188: 2465–2479 e2414

7. Burmann F, Lowe J (2023) Structural biology of SMC complexes across the tree of life. Curr Opin Struct Biol 80: 102598

8. Cutts EE, Tetiker D, Kim E, Aragon L (2025) Molecular mechanism of condensin I activation by KIF4A. EMBO J 44: 682–704

9. Cuylen S, Metz J, Haering CH (2011) Condensin structures chromosomal DNA through topological links. Nat Struct Mol Biol 18: 894–901

10. D’Ambrosio C, Schmidt CK, Katou Y, Kelly G, Itoh T, Shirahige K, Uhlmann F (2008) Identification of cis-acting sites for condensin loading onto budding yeast chromosomes. Genes Dev 22: 2215–2227

11. Davidson IF, Bauer B, Goetz D, Tang W, Wutz G, Peters JM (2019) DNA loop extrusion by human cohesin. Science 366: 1338–1345

12. Ganji M, Shaltiel IA, Bisht S, Kim E, Kalichava A, Haering CH, Dekker C (2018) Real-time imaging of DNA loop extrusion by condensin. Science 360: 102–105

13. Garcia-Rodriguez LJ, Kasciukovic T, Denninger V, Tanaka TU (2019) Aurora B-INCENP Localization at Centromeres/Inner Kinetochores Is Required for Chromosome Bi-orientation in Budding Yeast. Curr Biol 29: 1536–1544 e1534

14. Gibcus JH, Samejima K, Goloborodko A, Samejima I, Naumova N, Nuebler J, Kanemaki MT, Xie L, Paulson JR, Earnshaw WC et al (2018) A pathway for mitotic chromosome formation. Science 359

15. Giet R, Glover DM (2001) Drosophila aurora B kinase is required for histone H3 phosphorylation and condensin recruitment during chromosome condensation and to organize the central spindle during cytokinesis. J Cell Biol 152: 669–682

16. Griese JJ, Witte G, Hopfner KP (2010) Structure and DNA binding activity of the mouse condensin hinge domain highlight common and diverse features of SMC proteins. Nucleic Acids Res 38: 3454–3465

17. Hagstrom KA, Holmes VF, Cozzarelli NR, Meyer BJ (2002) C. elegans condensin promotes mitotic chromosome architecture, centromere organization, and sister chromatid segregation during mitosis and meiosis. Genes Dev 16: 729–742

18. Hirano M, Anderson DE, Erickson HP, Hirano T (2001) Bimodal activation of SMC ATPase by intra- and inter-molecular interactions. EMBO J 20: 3238–3250

19. Hirano M, Hirano T (1998) ATP-dependent aggregation of single-stranded DNA by a bacterial SMC homodimer. EMBO J 17: 7139–7148

20. Hirano M, Hirano T (2002) Hinge-mediated dimerization of SMC protein is essential for its dynamic interaction with DNA. EMBO J 21: 5733–5744

21. Hirano M, Hirano T (2006) Opening closed arms: long-distance activation of SMC ATPase by hinge-DNA interactions. Mol Cell 21: 175–186

22. Hirano T (2012) Condensins: universal organizers of chromosomes with diverse functions. Genes Dev 26: 1659–1678

23. Hu B, Itoh T, Mishra A, Katoh Y, Chan KL, Upcher W, Godlee C, Roig MB, Shirahige K, Nasmyth K (2011) ATP hydrolysis is required for relocating cohesin from sites occupied by its Scc2/4 loading complex. Curr Biol 21: 12–24

24. Kagami Y, Nihira K, Wada S, Ono M, Honda M, Yoshida K (2014) Mps1 phosphorylation of condensin II controls chromosome condensation at the onset of mitosis. J Cell Biol 205: 781–790

25. Kimura K, Hirano M, Kobayashi R, Hirano T (1998) Phosphorylation and activation of 13S condensin by Cdc2 in vitro. Science 282: 487–490

26. Kschonsak M, Merkel F, Bisht S, Metz J, Rybin V, Hassler M, Haering CH (2017) Structural Basis for a Safety-Belt Mechanism That Anchors Condensin to Chromosomes. Cell 171: 588–600 e524

27. Lammens A, Schele A, Hopfner KP (2004) Structural biochemistry of ATP-driven dimerization and DNA-stimulated activation of SMC ATPases. Curr Biol 14: 1778–1782

28. Lamothe R, Costantino L, Koshland DE (2020) The spatial regulation of condensin activity in chromosome condensation. Genes Dev 34: 819–831

29. Lavoie BD, Hogan E, Koshland D (2002) In vivo dissection of the chromosome condensation machinery: reversibility of condensation distinguishes contributions of condensin and cohesin. J Cell Biol 156: 805–815

30. Lavoie BD, Hogan E, Koshland D (2004) In vivo requirements for rDNA chromosome condensation reveal two cell-cycle-regulated pathways for mitotic chromosome folding. Genes Dev 18: 76–87

31. Lavoie BD, Tuffo KM, Oh S, Koshland D, Holm C (2000) Mitotic chromosome condensation requires Brn1p, the yeast homologue of Barren. Mol Biol Cell 11: 1293–1304

32. Lee BG, Merkel F, Allegretti M, Hassler M, Cawood C, Lecomte L, O’Reilly FJ, Sinn LR, Gutierrez-Escribano P, Kschonsak M et al (2020) Cryo-EM structures of holo condensin reveal a subunit flip-flop mechanism. Nat Struct Mol Biol 27: 743–751

33. Lipp JJ, Hirota T, Poser I, Peters JM (2007) Aurora B controls the association of condensin I but not condensin II with mitotic chromosomes. J Cell Sci 120: 1245–1255

34. Minnen A, Burmann F, Wilhelm L, Anchimiuk A, Diebold-Durand ML, Gruber S (2016) Control of Smc Coiled Coil Architecture by the ATPase Heads Facilitates Targeting to Chromosomal ParB/parS and Release onto Flanking DNA. Cell Rep 14: 2003–2016

35. Mishra A, Hu B, Kurze A, Beckouet F, Farcas AM, Dixon SE, Katou Y, Khalid S, Shirahige K, Nasmyth K (2010) Both interaction surfaces within cohesin’s hinge domain are essential for its stable chromosomal association. Curr Biol 20: 279–289

36. Mishra PK, Ciftci-Yilmaz S, Reynolds D, Au WC, Boeckmann L, Dittman LE, Jowhar Z, Pachpor T, Yeh E, Baker RE et al (2016) Polo kinase Cdc5 associates with centromeres to facilitate the removal of centromeric cohesin during mitosis. Mol Biol Cell 27: 2286–2300

37. Murayama Y, Uhlmann F (2015) DNA Entry into and Exit out of the Cohesin Ring by an Interlocking Gate Mechanism. Cell 163: 1628–1640

38. Nakazawa N, Mehrotra R, Ebe M, Yanagida M (2011) Condensin phosphorylated by the Aurora-B-like kinase Ark1 is continuously required until telophase in a mode distinct from Top2. J Cell Sci 124: 1795–1807

39. Petela NJ, Gligoris TG, Metson J, Lee BG, Voulgaris M, Hu B, Kikuchi S, Chapard C, Chen W, Rajendra E et al (2018) Scc2 Is a Potent Activator of Cohesin’s ATPase that Promotes Loading by Binding Scc1 without Pds5. Mol Cell 70: 1134–1148 e1137

40. Petersen J, Hagan IM (2003) S. pombe aurora kinase/survivin is required for chromosome condensation and the spindle checkpoint attachment response. Curr Biol 13: 590–597

41. Piazza I, Rutkowska A, Ori A, Walczak M, Metz J, Pelechano V, Beck M, Haering CH (2014) Association of condensin with chromosomes depends on DNA binding by its HEAT-repeat subunits. Nat Struct Mol Biol 21: 560–568

42. Pradhan B, Deep A, Konig J, Baaske MD, Corbett KD, Kim E (2025) Loop-extrusion-mediated plasmid DNA cleavage by the bacterial SMC Wadjet complex. Mol Cell 85: 107–116 e105

43. Pradhan B, Kanno T, Umeda Igarashi M, Loke MS, Baaske MD, Wong JSK, Jeppsson K, Bjorkegren C, Kim E (2023) The Smc5/6 complex is a DNA loop-extruding motor. Nature 616: 843–848

44. Rao SSP, Huang SC, Glenn St Hilaire B, Engreitz JM, Perez EM, Kieffer-Kwon KR, Sanborn AL, Johnstone SE, Bascom GD, Bochkov ID et al (2017) Cohesin Loss Eliminates All Loop Domains. Cell 171: 305–320 e324

45. Robellet X, Thattikota Y, Wang F, Wee TL, Pascariu M, Shankar S, Bonneil E, Brown CM, D’Amours D (2015) A high-sensitivity phospho-switch triggered by Cdk1 governs chromosome morphogenesis during cell division. Genes Dev 29: 426–439

46. Roisne-Hamelin F, Liu HW, Marechal N, Uchikawa E, Durand A, Gruber S (2025) Mechanism of DNA entrapment by a loop-extruding Wadjet SMC motor. Mol Cell 85: 3898–3912 e3897

47. Rojowska A, Lammens K, Seifert FU, Direnberger C, Feldmann H, Hopfner KP (2014) Structure of the Rad50 DNA double-strand break repair protein in complex with DNA. EMBO J 33: 2847–2859

48. Schalbetter SA, Goloborodko A, Fudenberg G, Belton JM, Miles C, Yu M, Dekker J, Mirny L, Baxter J (2017) SMC complexes differentially compact mitotic chromosomes according to genomic context. Nat Cell Biol 19: 1071–1080

49. Schwarzer W, Abdennur N, Goloborodko A, Pekowska A, Fudenberg G, Loe-Mie Y, Fonseca NA, Huber W, Haering CH, Mirny L et al (2017) Two independent modes of chromatin organization revealed by cohesin removal. Nature 551: 51–56

50. Soh YM, Burmann F, Shin HC, Oda T, Jin KS, Toseland CP, Kim C, Lee H, Kim SJ, Kong MS et al (2015) Molecular basis for SMC rod formation and its dissolution upon DNA binding. Mol Cell 57: 290–303

51. Srinivasan M, Scheinost JC, Petela NJ, Gligoris TG, Wissler M, Ogushi S, Collier JE, Voulgaris M, Kurze A, Chan KL et al (2018) The Cohesin Ring Uses Its Hinge to Organize DNA Using Non-topological as well as Topological Mechanisms. Cell 173: 1508–1519 e1518

52. St-Pierre J, Douziech M, Bazile F, Pascariu M, Bonneil E, Sauve V, Ratsima H, D’Amours D (2009) Polo kinase regulates mitotic chromosome condensation by hyperactivation of condensin DNA supercoiling activity. Mol Cell 34: 416–426

53. Sutani T, Yuasa T, Tomonaga T, Dohmae N, Takio K, Yanagida M (1999) Fission yeast condensin complex: essential roles of non-SMC subunits for condensation and Cdc2 phosphorylation of Cut3/SMC4. Genes Dev 13: 2271–2283

54. Tada K, Susumu H, Sakuno T, Watanabe Y (2011) Condensin association with histone H2A shapes mitotic chromosomes. Nature 474: 477–483

55. Tanaka S, Diffley JF (2002) Interdependent nuclear accumulation of budding yeast Cdt1 and Mcm2-7 during G1 phase. Nat Cell Biol 4: 198–207

56. Tane S, Shintomi K, Kinoshita K, Tsubota Y, Yoshida MM, Nishiyama T, Hirano T (2022) Cell cycle-specific loading of condensin I is regulated by the N-terminal tail of its kleisin subunit. Elife 11

57. Thadani R, Kamenz J, Heeger S, Munoz S, Uhlmann F (2018) Cell-Cycle Regulation of Dynamic Chromosome Association of the Condensin Complex. Cell Rep 23: 2308–2317

58. Uhlmann F (2016) SMC complexes: from DNA to chromosomes. Nat Rev Mol Cell Biol 17: 399–412

59. Verzijlbergen KF, Nerusheva OO, Kelly D, Kerr A, Clift D, de Lima Alves F, Rappsilber J, Marston AL (2014) Shugoshin biases chromosomes for biorientation through condensin recruitment to the pericentromere. Elife 3: e01374

60. Wells JN, Gligoris TG, Nasmyth KA, Marsh JA (2017) Evolution of condensin and cohesin complexes driven by replacement of Kite by Hawk proteins. Curr Biol 27: R17–R18

61. Wilhelm L, Burmann F, Minnen A, Shin HC, Toseland CP, Oh BH, Gruber S (2015) SMC condensin entraps chromosomal DNA by an ATP hydrolysis dependent loading mechanism in Bacillus subtilis. Elife 4

62. Woo JS, Lim JH, Shin HC, Suh MK, Ku B, Lee KH, Joo K, Robinson H, Lee J, Park SY et al (2009) Structural studies of a bacterial condensin complex reveal ATP-dependent disruption of intersubunit interactions. Cell 136: 85–96

